# Putative azithromycin-resistance mutations in *Chlamydia trachomatis* are globally distributed but arose before azithromycin was discovered

**DOI:** 10.1101/2025.11.10.687758

**Authors:** Parul Sharma, Deborah Dean, Timothy D. Read

**Author notes:** **Corresponding authors:** Deborah Dean, Timothy D. Read.

## Abstract

Azithromycin is widely used to treat *Chlamydia trachomatis* infections, yet the extent of resistance to the drug across the species has not been addressed. We surveyed mutations and substitutions linked to putative azithromycin resistance across 1,354 high-quality *C. trachomatis* genomes. Mutations in the *rpl*V gene encoding three non-synonymous substitutions, compared to the canonical *C. trachomatis* reference strain D//TW-3/Cx sequence, were found to be common but largely conserved within lineages causing prevalent urogenital and anorectal infections and lymphogranuloma venereum (LGV). Time-scaled phylogenetic analysis suggested that these mutations predate the clinical introduction of azithromycin. In contrast, no consistent resistance-associated patterns were observed in *23S* rRNA or *rpl*D genes. This large-scale genomic surveillance provides critical insights into the evolutionary trends of putative azithromycin resistance in *C. trachomatis* and underscores the importance of integrating genomic monitoring with phenotypic susceptibility testing to accurately assess and manage antimicrobial resistance.

## Introduction

*Chlamydia trachomatis* is an obligate intracellular pathogen and the most common cause of bacterial sexually transmitted infections (STIs) worldwide. Ocular strains of *C. trachomatis* are also the leading cause of infectious blindness, particularly among women. In STIs, treatment commonly involves antibiotics such as doxycycline administered twice a day over seven days or azithromycin as a single dose (“Chlamydia,” n.d.). However, in trachoma-endemic regions, topical tetracycline ointment or oral azithromycin have historically been the preferred options (Dawson et al. 1997). The convenience of a single-dose regimen has made azithromycin a preferred option, supporting higher treatment adherence. It is the antibiotic of choice in many mass drug administration (MDA) campaigns implemented under the World Health Organization (WHO) SAFE (Surgery, Antibiotics, Facial cleanliness, and Environmental improvement) strategy, aimed at eliminating blinding trachoma ((“WHO Guideline on Mass Drug Administration of Azithromycin to Children under Five Years of Age to Promote Child Survival,” n.d.). Despite large-scale efforts, elimination targets have not been fully achieved, and several countries, particularly in sub-Saharan Africa such as Ethiopia and Sudan, remain endemic. In recognition of these challenges, the WHO extended its global elimination goal for blinding trachoma from 2020 to 2030 (O’Brien et al. 2019; Ageed and Khan 2024).

Azithromycin is an azalide, a sub-class of macrolide antibiotics (Bakheit, Al-Hadiya, and Abd-Elgalil 2014). It was discovered in 1980, patented in 1981, and its widespread use began after 1991 when it was launched by Pfizer (Tomišić2011). Since then, it has become one of the most widely prescribed antibiotics for bacterial infections, including *C. trachomatis*, due to its broad-spectrum activity, tissue penetration and long half-life, and patient compliance advantages (Ballow and Amsden 1992; Zhang et al. 2025). However, its extensive use has also raised concerns about the potential for emerging resistance .

Several STI studies have reported *C. trachomatis* treatment failures with azithromycin (Peterman et al. 2006; Kissinger et al. 2016; Batteiger et al. 2010). These failures are often attributed to either reinfection of another *C. trachomatis* strain after clearance of the initial infection or persistence of the initial infection. Persistent infections have been attributed to development of *C. trachomatis* antibiotic resistance (Batteiger et al. 2010; Somani et al. 2000). Several studies have identified point mutations in 23S rRNA, *rplV* and *rplD* genes as potentially linked to macrolide resistance in *C. trachomatis* (Rachel Binet and Maurelli 2007; Benamri et al. 2021; Lodhia et al. 2025). In *rpl*V, encoding the L22 protein, mutations resulting in G52S, R65C, and V77A substitutions were detected in isolates from patients with azithromycin treatment failure (Deguchi et al. 2018; Zhu et al. 2010), although the minimum inhibitory concentrations (MIC) were within the sensitive range. The same variants were observed in Russian urogenital isolates, where they were present in both macrolide-resistant and sensitive strains (Misyurina et al. 2004), and among endocervical, vaginal and rectal samples in Fiji (Bommana et al. 2025).

For RplD, the L4 protein, studies reported P109L, P151A (Zhu et al. 2010) and Q66K substitutions (Rachel Binet and Maurelli 2007), while other studies (Bhengraj, Srivastava, and Mittal 2011; Deguchi et al. 2018) did not detect *rpl*D mutations in clinical isolates. 23S rRNA gene mutations A2057G, A2058C, A2059G, and T2611C, have been associated with resistance, although their presence and role remain inconsistent across studies (Jiang et al. 2015; Misyurina et al. 2004; Xue et al. 2017; Zhu et al. 2010). Notably, all the above 23S rRNA mutations are also listed in the NCBI Reference Gene Catalog (Feldgarden et al. 2021), a public database that compiles known antimicrobial resistance genes and associated mutations across diverse pathogens. Similarly, the Comprehensive Antibiotic Resistance Database (CARD) (Alcock et al. 2020) serves as another curated resource detailing resistance determinants, mechanisms, and mutation-based associations. The latest version of CARD (accessed Oct, 2025) specifically documents *23S* rRNA mutations linked to azithromycin resistance in *C. trachomatis*, while the NCBI catalog lists *rpl*D, *rpl*V, and *23S* rRNA mutations reported across multiple bacterial species.

In this study, we analyze the patterns of DNA mutations in the 23S rRNA gene and amino acid substitutions in L22 and L4 associated with azithromycin resistance across a globally distributed collection of 1,354 *C. trachomatis* genomes from public databases. Our aim was to characterize the prevalence, lineage specificity, and evolutionary patterns of putative azithromycin resistance variants. Since antimicrobial resistance in *C. trachmatis* remains significantly understudied, we also scanned for novel variants that had a pattern of convergent evolution on the phylogeny. The independent appearance of the same mutation could be suggestive of emergence of resistance under the selective pressure of azithromycin (Card et al. 2021).

## Methods

### Compiling and pre-processing genomic data

*C. trachomatis* genomes were compiled from multiple sources, including the NCBI Assembly database (accessed March, 2025), AllTheBacteria database (version 0.2 with the incremental release from 08-2024) (Hunt et al. 2024), published reference genomes (Olagoke et al. 2025), and available NCBI SRA samples. For NCBI SRA samples, raw-reads were down-sampled to achieve even coverage using the bbnorm.sh script from BBMap (v39.01) (Bushnell 2014), followed by assembly with SPAdes (v4.0.0) (Prjibelski et al. 2020) using default parameters. Quality filtration was applied across the complete dataset of genomes from all sources using checkM (v1.2.3) (Parks et al. 2015) retaining only genomes with more than 98% completeness, less than 6% contamination and less than 2000 contigs.

### Identifying azithromycin resistance-associated mutations

A list of point mutations and substitutions linked to macrolide resistance was obtained from the NCBI Reference Gene Catalog (accessed March, 2025) (Feldgarden et al. 2021) and CARD (Alcock et al. 2020). Additional candidate mutations reported in the literature for azithromycin resistance in *C. trachomatis* were also compiled (**Table 1**). To ensure comparability, all mutations were converted to *C. trachomatis* numbering by alignment to *C. trachomatis* reference strain D/UW-3/Cx (NCBI Accession ID: NC_000117.1) (Stephens et al. 1998).

**Table 1.** List of azithromycin associated 23S mutations in public databases.

| Source | Allele | Reference | Reference species | CT-numering | Detected |
| --- | --- | --- | --- | --- | --- |
| NCBI | a2074c /<br>a2074g /<br>a2074t | NC_022347.1,<br>NC_002163.1 | *Campylobacter coli*,<br>*Campylobacter jejuni* | a1879c /<br>a1879g /<br>a1879t | yes (a->g) |
|  | a2075g | NC_022347.1,<br>NC_002163.1 | *Campylobacter coli*,<br>*Campylobacter jejuni* | a1880g |  |
|  | a2059g | NC_002946.2 | *Neisseria gonorrhoeae* | a2036g | yes |
|  | a2058g /<br>a2058t | NC_004431.1 | *Escherichia coli* | a2038g |  |
|  | a2114g | NZ_CP018347.1 | *Streptococcus pneumoniae* | a2449g |  |
|  | a2115g | NZ_CP018347.1 | *Streptococcus pneumoniae* | a2450g |  |
|  | c2627a | NC_022347.1,<br>NC_002163.1 | *Campylobacter coli*,<br>*Campylobacter jejuni* | c2592a |  |
|  | c2611t | NC_002946.2,<br>NC_004431.1 | *Neisseria gonorrhoeae*,<br>*Escherichia coli* | c2592t /<br>c2592g | yes (c->t) |
|  | g2032t | NC_004431.1 | *Escherichia coli* | g2012t |  |
|  | c2630a /<br>c2630g | NZ_CP018347.1 | *Streptococcus pneumoniae* | na* |  |
|  | t2609c | NC_004431.1 | *Escherichia coli* | t2590c |  |
|  | t754a | NC_004431.1 | *Escherichia coli* | t740a | yes (t->g) |
| CARD | c196a | NR_076160.1 | *Chlamydia trachomatis* | c196a |  |
|  | a2107c /<br>a2107g | NR_076160.1 | *Chlamydia trachomatis* | a2107c /<br>a2107g |  |
|  | a2109g | NR_076160.1 | *Chlamydia trachomatis* | a2109g |  |
|  | a2663c /<br>t2663c | NR_076160.1 | *Chlamydia trachomatis* | a2663c /<br>t2663c |  |

\*na = not aligned. Detected mutations were present in more than 99% of the isolates. *C. trachomatis*-numbering was determined by aligning the reference sequence against *C. trachomatis* reference NC\_000117.1

All genomes in the dataset were re-annotated using Bakta (v1.9.2) (Schwengers et al. 2021) to minimize annotation heterogeneity, and sequences corresponding to the 23S rRNA, *rpl*V, and *rpl*D genes were extracted. Each gene set was aligned to the reference sequence using MAFFT (v7.526) (Katoh and Standley 2013), and variants were identified relative to the reference coordinates. Mutations were detected and summarized using custom python scripts (available on GitHub:https://github.com/parul-sharma/CT-AMR).

### Phylogenetic trees and time-scaled analysis

Core-genome genes were identified for all annotated genomes in the dataset using PIRATE (v.1.0.5) (Bayliss et al. 2019) under a single workflow. The resulting core gene alignment, composed of 853 genes 0.83Mb in length, was used as input for IQ-TREE2 (version 2.3.0) (Minh et al. 2020) to construct a maximum-likelihood phylogenetic tree with automated model selection. A similar phylogenetic tree was obtained for the subset of genomes (138 out of 1354) with associated dates of isolation. For these dated strains, dates of isolation were added using the ‘–date’ parameter to supply temporal metadata to the IQ-TREE run. The resulting tree was subsequently used as input for BEAST (v2.7.7) (Bouckaert et al. 2019) to perform time-scaled phylogenetic inference.

Using BEAST, we evaluated multiple models for phylogenetic inference. All models used a strict molecular clock but differed in population models: (i) Bayesian Skyline; (ii) Bayesian Skyline with HKY nucleotide substitution; (iii) exponential growth; (iv) constant population size; and (v) extended Bayesian Skyline with invariant sites. All models produced broadly similar topologies and divergence times (**Supplementary Data**). For clarity, the results presented here are based on the strict molecular clock with an exponential growth population model. Random Local Clock models were not attempted, as the low overall divergence of *C. trachomatis* sequences would provide insufficient signal to reliably estimate lineage-specific rate variation.

### Recombination and visualizations

Recombination analysis was performed on the core genome alignment (described above) using ClonalFrameML (v1.13) (Didelot and Wilson 2015) with default parameters. The resulting importation status file was processed with a custom python script (available on GitHub https://github.com/parul-sharma/CT-AMR) to identify recombination regions. Briefly, the genome was divided into 1,000 bp windows, the total number of recombination events within each window was counted, and windows with elevated recombination activity (more than 20 recombination events) were designated as recombination “hotspots”.

To investigate whether resistance-associated loci showed phylogenetic patterns distinct from the core genome, we constructed individual gene trees for the 23S, *rpl*V, and *rpl*D sequences. These were then compared to the core-genome phylogeny using tanglegrams, allowing us to check congruence in tree structure and evaluate potential evidence of recombination or lineage-specific inheritance at these loci. All visualizations were generated in R, with trees constructed using ggtree (v3.13.0) (Xu et al. 2022), time-scaled trees in combination with treeio (v1.28.0) (Wang et al. 2020), and tanglegram comparisons performed using phytools (v2.4-4) (Revell 2012). Supporting R scripts are available in the GitHub repository.

## Results

### Putative-resistance mutations in the *rplV* gene are lineage specific

We examined resistance-associated mutations in our dataset of 1,354 high-quality *C. trachomatis* genomes (**Supplementary Table 1**). A core-genome phylogeny divided the genomes into four major lineages, consistent with prior studies: lymphogranuloma venereum (LGV), ocular, prevalent urogenital/anorectal (P-UA), and non-prevalent urogenital/anorectal (NP-UA), reflecting their disease tropisms (Joseph et al. 2023; Harris et al. 2012; Joseph et al. 2012). All substitutions were called in relation to the canonical reference D/UW-3/Cx sequence (NC_000117.1), an NP-UA genome (Stephens et al. 1998).

We first focused on the three amino acid substitutions in L22: G52S, R65C, and V77A, that have been associated with azithromycin resistance in *C. trachomatis* (Misyurina et al. 2004; Deguchi et al. 2018). In each case, we found that the substitution was caused by a single mutation (g151a, c190t and t227c, respectively). These substitutions showed strong lineage specificity (**Fig1**). All LGV genomes (193/193, 100%) carried G52S and V77A, while nearly all P-UA genomes (399/404, 98.8%) harbored the complete set of triple substitutions (G52S, R65C, and V77A). In contrast, these putative resistance-associated substitutions were completely absent from ocular genomes (0/451) and were rare in NP-UA genomes (4/306, 1.3%). In the rare NP-UA genomes with the G52S, R65C, and V77A substitutions, the sequence was identical to the P-UA allele, suggesting exchange by homologous recombination. A similar pattern in reverse was seen in the rare P-UA genomes missing these mutations (**Supplementary Table 2**).

**Table 2.**
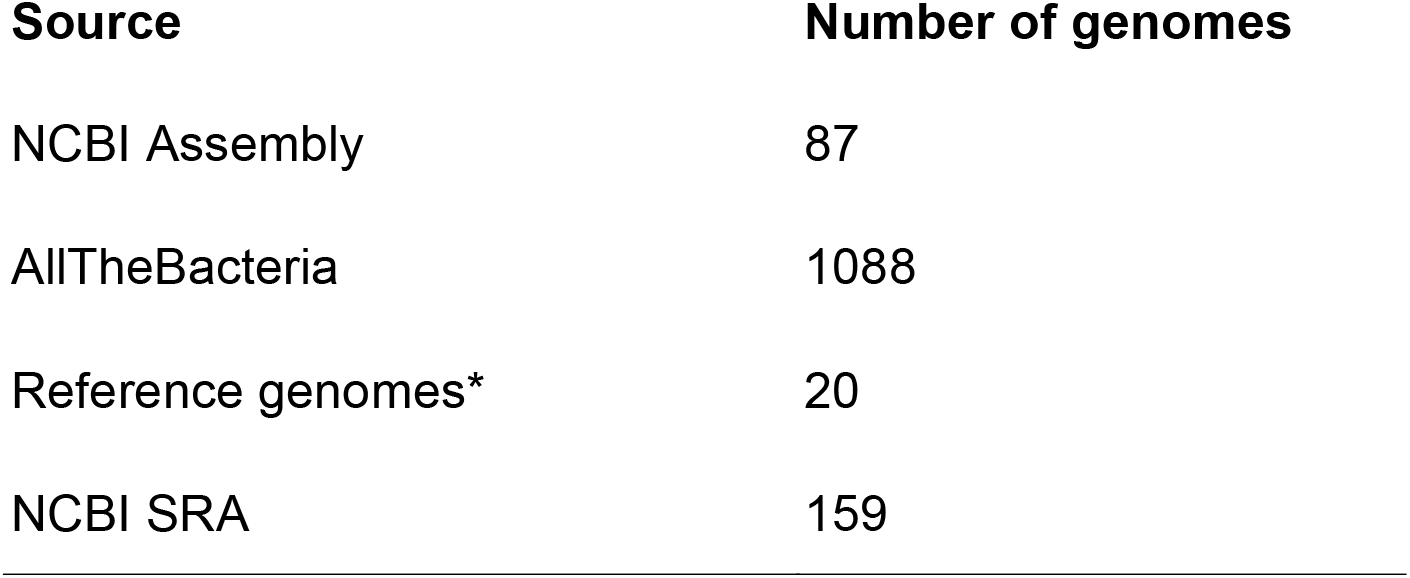
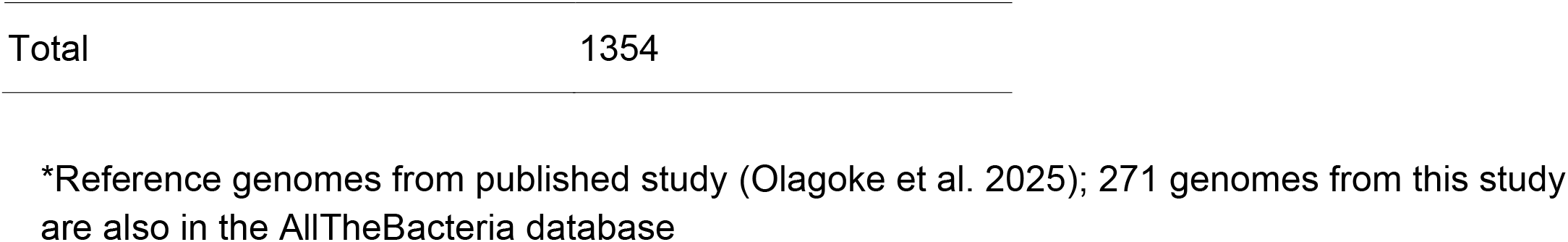
Number of genomes derived from each data source.

We next broadened our analysis to identify additional *rpl*V mutations that resulted in amino acid substitutions relative to the reference D/UW-3/Cx protein sequence (accession ID: NC_000117.1) across the dataset. Thirteen other substitutions were detected, most at low frequencies and restricted to specific lineages with the only exception being the Q111R substitution present in few of both Ocular (28/451, 6.2%) and LGV (2/193, 1.0%) genomes (**Fig1**). In the P-UA lineage, S86A and K87R co-occurred in a single genome, while L22S was detected in another. Among ocular genomes, R17H was the most common secondary variant (86/451, 19.1%), followed by Q111R (28/451, 6.2%), with several others present at very low frequencies (N25D, E30K, and E108G, each in ≤2 genomes). In NP-UA genomes, V29I was observed in 26/306 (8.5%), I63V in 4/306 (1.3%), and T99S in a single genome. Within LGV genomes, Q111R was present in 2/193 (1.0%), while V70I was more frequent (69/193, 35.8%); R65H occurred only once.

Together, these findings demonstrated that the putative azithromycin-associated *rplV* mutations were tightly clustered in P-UA and LGV lineages. There were other substitutions but, aside from Q111R, there was no evidence of homoplasy. There was also evidence of gene conversion events in the P-UA and NP-UA lineage strains.

### L22 resistance-associated substitutions predate the widespread clinical use of azithromycin

For a subset of 138 (10.2%) of the 1354 genomes, metadata on the year of isolation was available (**Supplementary Table 1**). Genome SAMEA767935 (P-UA lineage, E/Bour ompA type), collected in 1959 (Hanna, Thygeson, and Jawetz 1959), carried all three canonical L22 resistance-associated substitutions (G52S, R65C, and V77A). Another 1959 genome, SAMEA1973344 (Ocular lineage, C/TW-3/OT ompA type), had no substitutions in L22. The second-oldest genome, SAMEA767923 (LGV lineage, L3/404 ompA type), collected in 1967, carried the double mutations (G52S and V77A).

Azithromycin was first discovered in 1981 and approved for clinical use after 1991 (Tomišić 2011). The presence of resistance-associated *rplV* mutations in genomes dating from 1959 and 1967 indicated that these mutations arose prior to the widespread clinical use of this antibiotic. Time-scaled phylogenetic analysis (**Fig. 2**) further supported this observation. Using the strict molecular clock with an exponential growth population model, we estimated a 95% likelihood that genomes carrying the triple mutations diverged between 1016–1779, while those with the double mutations diverged between 850–1695.

**Fig 1.**
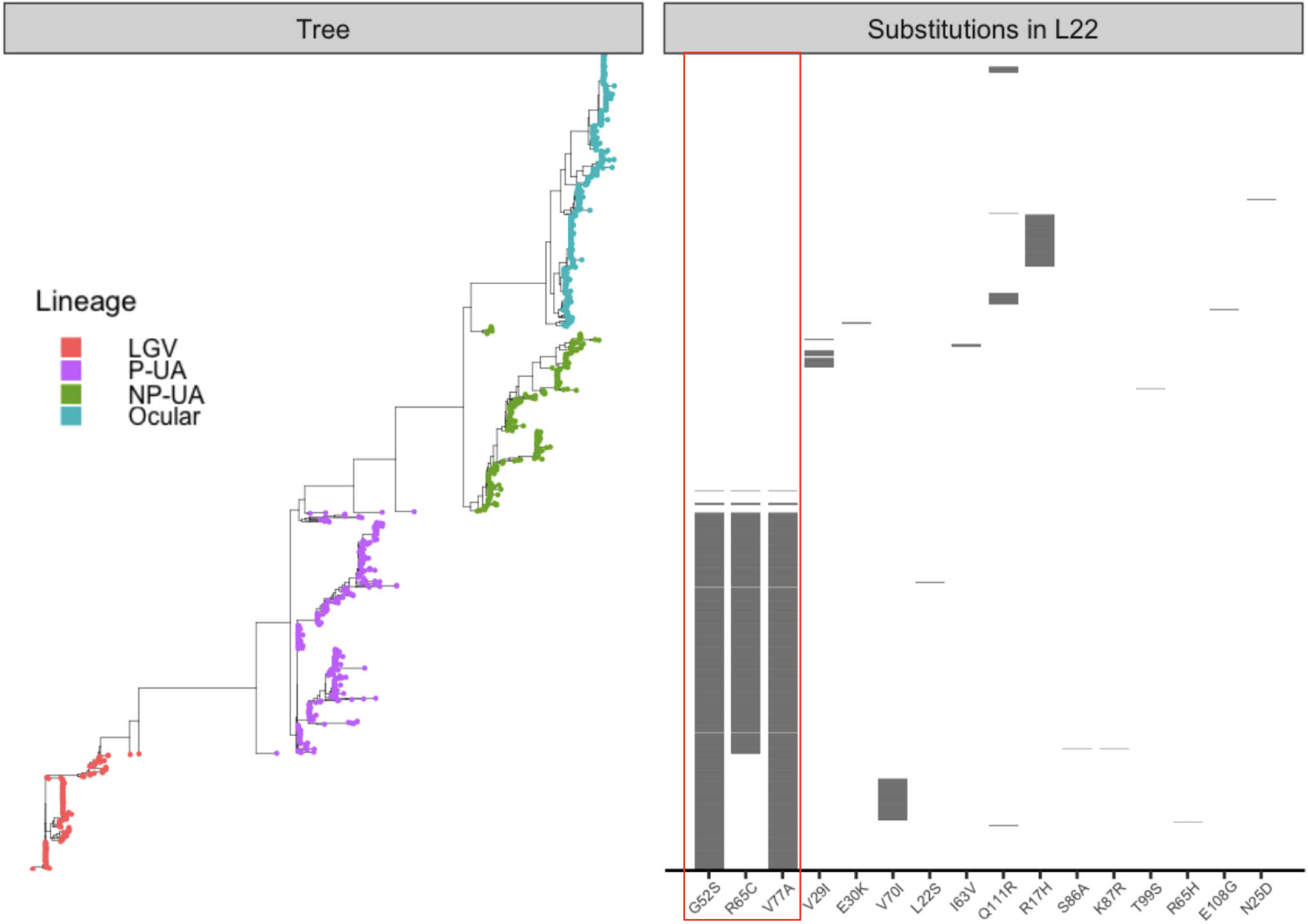
Distribution of L22 substitutions across genomes. Genomes are ordered according to core-gene phylogeny (left) with tips colored by *C. trachomatis* lineage (LGV, ocular, P-UA, NP-UA). The panel on the right shows a heatmap of the 16 L22 substitutions, marked along the x-axis, with grey tiles indicating presence and empty spaces indicating absence across samples. The three azithromycin-associated substitutions are marked with a red box.

**Fig 2.**
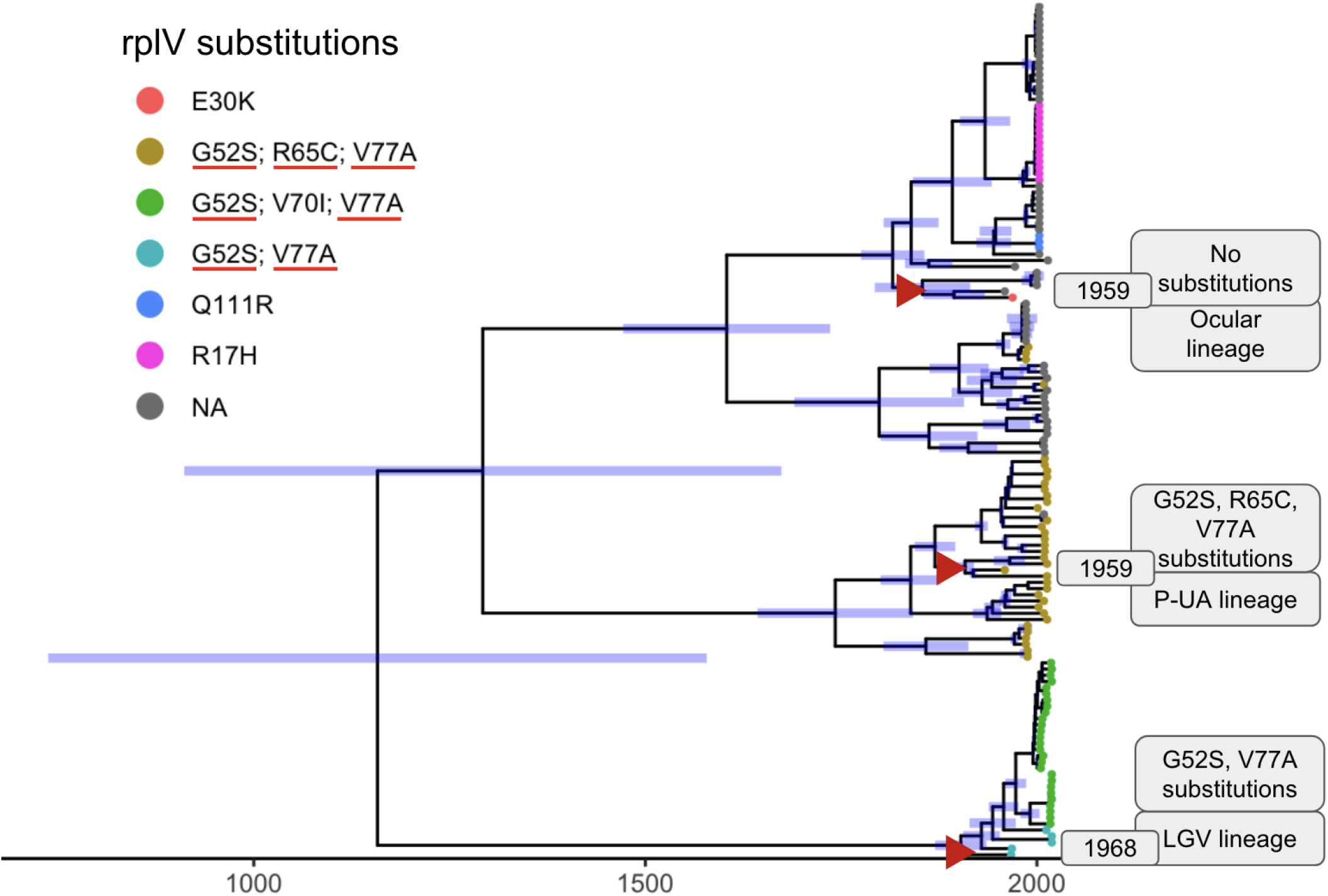
Time-scaled phylogeny constructed using BEAST. Branch lengths are proportional to time (in years), and the 95% highest posterior density (HPD) intervals of node heights are indicated as blue bars. Tip labels annotated by color representing **rplV** substitutions as per the legend with azithromycin-associated substitutions marked with red underline. The three oldest genomes (highlighted with red triangles) are annotated with metadata including year of isolation, lineage, and presence of **rplV** substitutions.

### Patterns of mutations in *rpl*D and 23S rRNA

The three L4 protein substitutions previously reported as azithromycin resistance-associated (P109L, P151A (Zhu et al. 2010) and Q66K (R. Binet and Maurelli, n.d.)) were not found across the 1354 *C. trachomatis* genomes in this study. Due to the high sequence divergence of the *rpl*D gene across bacterial species, none of the putative resistance-associated substitutions listed in the NCBI Reference Gene Catalog could be identified in our dataset, as protein sequences from other species did not align with the *C. trachomatis* L4 protein (**Supplementary Table 3**). A small number of substitutions were observed within the protein, most of which were lineage-specific (**Fig. 3**). Notably, two homoplasic substitutions - R111Q and V132I - were predominant in P-UA genomes (399/404, 98.8%), though they also appeared in a few NP-UA (n=2), Ocular (n=3), and LGV (n=6) genomes. These variants were absent in five P-UA genomes and partially missing in three others, missing R111Q in two and V132I in one (**Supplementary Table 4**).

**Fig 3.**
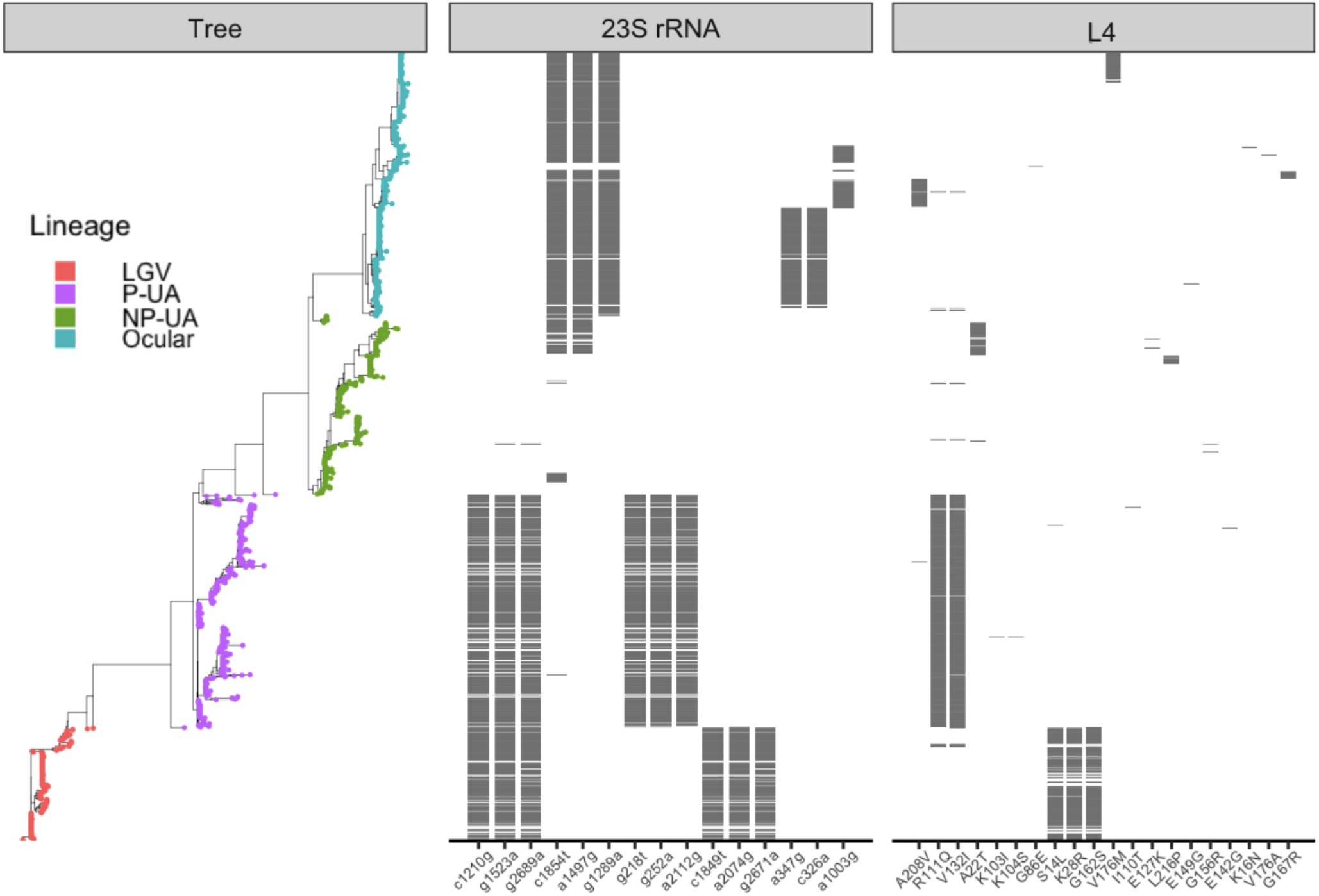
Distribution of mutations in 23S rRNA and substitutions in L4 across genomes. Genomes are ordered by core-gene phylogeny (left), with tree tips colored by genome lineage. The first panel shows the 15 most frequent 23S rRNA mutations; the second panel shows all L4 substitutions. In both panels, the x-axis lists individual mutations or substitutions.

Similarly, examination of the 23S rRNA sequence showed no obvious correlation with previously reported resistance mutations. Of the ten resistance-associated positions previously described in NCBI AMR and CARD databases (**Table 2**), four were present in all strains, including the reference strain used for alignment. This suggests that these variants represent fixed or highly conserved positions rather than markers of acquired resistance. Across the dataset, a total of 104 additional mutations were detected, ranging from 1 to 8 mutations per genome, with 77.8% (81 out of 104) present in fewer than 1% of genomes. Additionally, since *C. trachomatis* carries two copies of the 23S rRNA gene, we examined whether any mutations were heterozygous.

Only 11 of the 89 complete genome assemblies exhibited heterozygous mutations, all of which were rare (present in less than 1% of genomes). To better understand lineage-specific patterns, we therefore visualized the15 most frequent mutations. The 15 most prevalent mutations were non-homoplasious and largely lineage-specific, with genomes from the NP-UA lineage displaying the most rare mutations. However, the observed frequencies of mutations was influenced by the use of the NP-UA type strain (NC_000117.1, D/UW-3/Cx) as the reference for mutation calling, meaning that all differences were computed relative to this sequence.

Collectively, these observations indicate that, similar to *rpl*V, variation in *rpl*D and 23S rRNA is largely shaped by lineage and genomic background, and there is no strong evidence for recent selection for azithromycin resistance. While allelic diversity is largely lineage-dependent, there is no evidence that these loci have been the primary drivers of resistance emergence in *C. trachomatis*.

### *rpl*V, *rpl*D and 23S rRNA are not in genomic recombination hotspots

To investigate whether resistance-associated mutations in *rpl*V, *rpl*D, or 23S rRNA arose via intra-specific recombination, we compared gene-specific phylogenies with the core-genome phylogeny for a random subset of genomes. Across all comparisons, *rpl*V vs. core tree, *rpl*D vs. core tree, and 23S rRNA vs. core tree (**Supplementary Data**), we observed strong congruence, with no evidence of anomalous branching patterns indicative of horizontal transfer or recombination. The observation was further supported by genome-wide recombination analyses. None of the sequences in these ribosomal genes were located within identified recombination hotspots (**Figure 4**). In contrast, recombination events were predominantly observed in expected loci, such as genes encoding membrane proteins, with *ompA* exhibiting the highest density of recombination sites.

**Fig 4.**
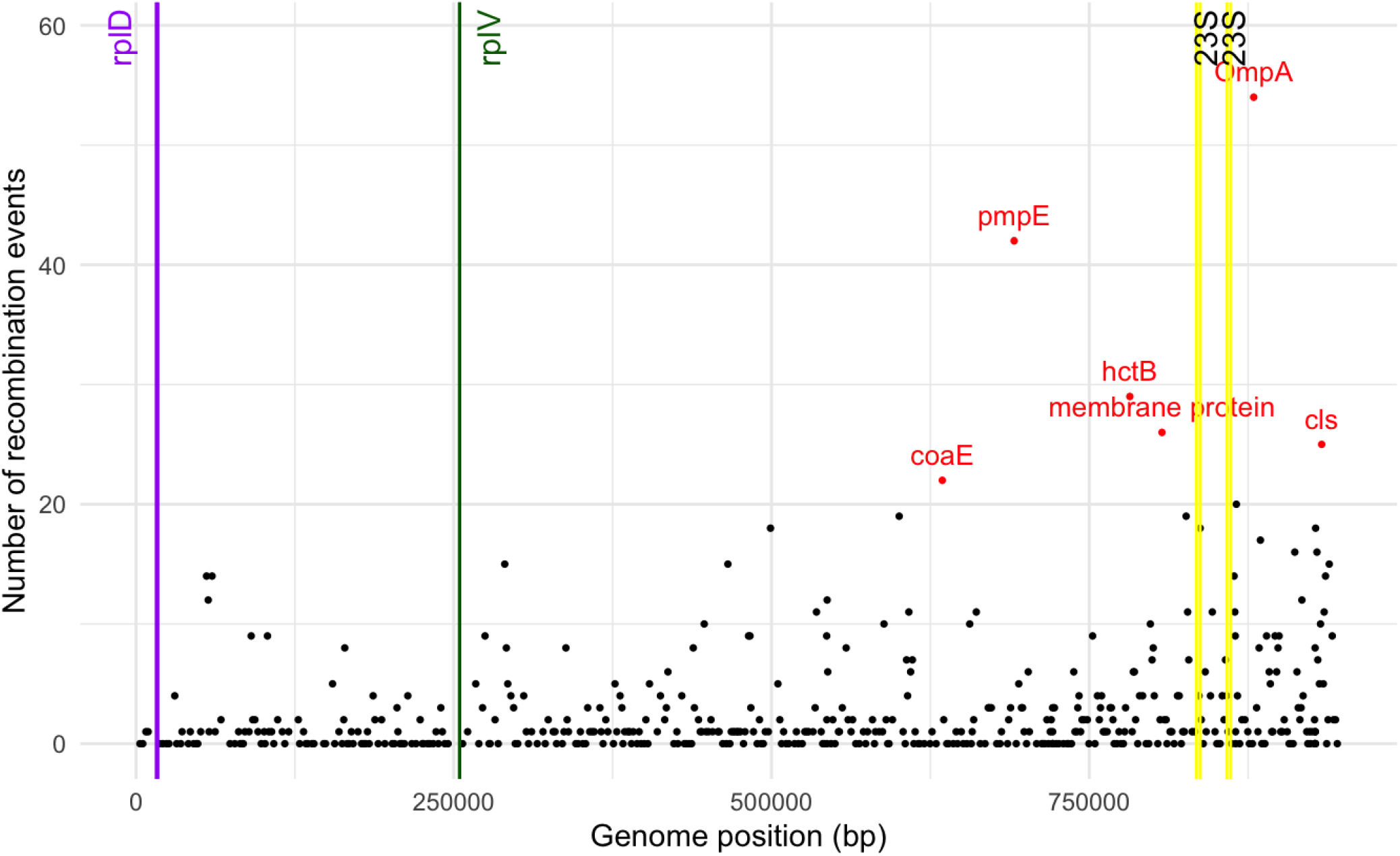
Recombination hotspots across *C. trachomatis* genome. Frequency of recombination events mapped to the *C. trachomatis* reference strain D/UW-3/Cx (NCBI Accession: NC_000117.1). The vertical lines indicate the genomic positions of the *rplD* (purple), *rplV* (green), and two copies of the 23S rRNA (yellow) genes. Regions with more than 20 recombination events are shown as red dots, representing recombination hotspots across the *C. trachomatis* genome.

## Discussion

Many methods of inference of resistance from genome sequences require catalogs of mutations linked to phenotypes (Su, Satola, and Read 2019). Here, using 1,354 available *C. trachomatis* genomes, we investigated the phylogenetic patterns of previously characterized mutations that were purported to be linked to azithromycin resistance, but varied in the degree of susceptibility or resistance based on MICs. The increasing availability of large public genomic datasets now provide an opportunity to systematically explore resistance-associated mutations across diverse populations with more comprehensive datasets. This large-scale genomic approach provides a framework for better understanding resistance dynamics in *C. trachomatis* and lays the foundation for improved surveillance and clinical management strategies.

Our study provides evidence that putative azithromycin resistance alleles were already present in *C. trachomatis* lineages before the widespread clinical use of azithromycin that occurred after 1991. The high lineage specificity of these alleles suggests that their persistence more likely reflects neutral evolution rather than resistance acquired after azithromycin use. These findings align with a growing body of work demonstrating that the genetic foundations of antimicrobial resistance often predate human antibiotic use (D’Costa et al. 2011), with ancient selective forces shaping the genomic background upon which modern resistance emerges (Kaul et al.

2025). However, the mutations may not be directly linked to elevated resistance to azithromycin. Possibly, the putative resistance phenotype linked to *rplV* found by other groups (Misyurina et al. 2004; Deguchi et al. 2018) is linked to additional factors such as epistatic mutations (Wong 2017). Nonetheless, these findings underscore the importance of considering historical and lineage-specific variation when interpreting the possible emergence of antibiotic resistance in *C. trachomatis*.

Pinpointing azithromycin resistance in *C. trachomatis* is complicated by several factors. Although treatment failures with azithromycin have been repeatedly reported, these events are multifactorial in nature. Reinfection from an untreated partner, inadequate drug treatment, or noncompliance with medication are common confounders that can mimic resistance-associated persistence (Kissinger et al. 2016; Hocking et al. 2013). Moreover, while genotypic evidence of resistance alleles provides valuable insights, it cannot be equated directly to a resistance phenotype. Several older studies highlighted this disconnect, where treatment failures were observed without clear links to resistance-associated mutations, underscoring the complexity of resistance as a trait (Shao et al. 2020). Such observations reinforce the idea that resistance in *C. trachomatis* is a multifactorial trait, with potential contributions from epistatic mutations, epigenetic regulation, host-pathogen interactions, or other yet-uncharacterized mechanisms.

Our analysis also underscores the limitations of transferring resistance markers across species. Variants annotated in public resistance databases, often curated based on other pathogens, do not consistently predict resistance in *C. trachomatis*. For example, macrolide resistance mutations in 23S rRNA (e.g., A2059G, A2074G, C2611T), which are well established in *Neisseria gonorrhoeae, Campylobacter coli*, and *Escherichia coli*, do not show consistent associations with resistance in *C. trachomatis*. In fact, we found these mutations in nearly all genomes in our dataset, indicating that they likely reflect natural variation rather than resistance. This lack of correlation highlights the non-transferability of resistance markers between species. Instead, 23S variation in *C. trachomatis* appears to track more closely with phylogenetic background than with resistance, underscoring the need for organism-specific surveillance.

Reliance on databases built primarily from other pathogens may otherwise overstate or misrepresent the clinical relevance of certain variants, especially when extrapolating resistance annotations across taxa.

Since antimicrobial resistance in *C. trachomatis* remains significantly understudied, we also looked for previously uncharacterized mutations in the three loci that may have an evolutionary pattern suggestive of association with resistance. We theorized that homoplasic mutations (i.e., independent appearance of the same mutation in separate lineages) would be a signal of selection pressure due to antibiotic treatment (Card et al. 2021). There was only one L22 substitution, Q111R, and two L4 substitutions, R111Q and V132I, that had homoplasious patterns. While nothing is known about the phenotypic effect of the underlying mutations, this phylogenetic pattern suggests the need for further investigation. Interestingly, there were also rare cases of apparent recombination swapping of *rpl*V alleles between P-UA and NP-UA genomes. However, the three loci (*rpl*V, *rpl*D and 23S rRNA) were not hotspots for recombinational transmission in general, which also might be expected of novel drug resistance mutations. Instead, mutations were most often rare and lineage dependent.

Major challenges in advancing *C. trachomatis* antimicrobial resistance research lies in the lack of phenotypic confirmation. Due to its obligate intracellular lifestyle, traditional culture-based MIC testing is technically challenging, labor-intensive, and rarely performed in routine laboratories (Elwell, Mirrashidi, and Engel 2016). This reliance on limited phenotypic data has constrained our ability to validate resistance mutations at scale. Nonetheless, in-depth genomic analyses such as ours are crucial for prioritizing candidate alleles and selecting samples for functional studies. By refining the pool of targets, these studies provide a framework to design phenotypic assays more efficiently, ultimately bridging the gap between genotypic surveillance and clinical relevance.

Taken together, our findings reinforce several key messages. First, resistance-associated alleles in *C. trachomatis* are shaped by lineage-specific evolution and likely predate clinical use of azithromycin. Second, mutations annotated as resistance markers in public databases derived from other pathogens cannot be assumed to apply to *C. trachomatis*. Third, while large-scale genomic surveys provide valuable insights into the evolutionary context of putative resistance mutations, they cannot on their own establish clinical relevance. Functional validation through phenotypic MIC testing and experimental work remains essential. Moving forward, integrating genomic surveillance with phenotypic evidence will be key to resolving the complex relationship between genetic variation and antimicrobial resistance in *C. trachomatis*, ensuring that resistance monitoring and treatment strategies are both accurate and tailored to this pathogen’s unique biology.

## Conclusion

This large-scale genomic survey of 1,354 *Chlamydia trachomatis* genomes revealed that mutations in the *rpl*V gene are abundant and strongly conserved within infectious lineages, particularly prevalent urogenital and rectal (P-UA) and LGV strains, whereas no consistent resistance-associated patterns were observed in the 23S rRNA or *rpl*D genes. These results indicate that known resistance mutations are not universal predictors of resistance in *C. trachomatis* and instead reflect lineage-specific evolutionary histories that predate the clinical use of azithromycin. Our findings highlight the limitations of directly transferring resistance markers from other pathogens and demonstrate that variation in *C. trachomatis* often tracks with phylogenetic background rather than resistance. Moving forward, organism-specific genomic surveillance, coupled with phenotypic validation, will be critical to disentangle the evolutionary and functional bases of resistance in *C. trachomatis*. Together, these insights provide a foundation for more accurate monitoring and treatment strategies tailored to the unique biology of this pathogen.

## Acknowledgement

This study was supported by the National Institutes of Health (NIH) grants AI51075 and AI158527 awarded to TDR and DD.

## List of figures and tables

^⋆^Reference genomes from published study (Olagoke et al. 2025); 271 genomes from this study are also in the AllTheBacteria database

Supplementary Data: (i) Comparisons of core-phylogenetic tree with gene trees (RplV, RplD and 23S rRNA), (ii) Comparisons of different BEAST models

Supplementary Table 1: Metadata of all high-quality samples used for the study

Supplementary Table2: List of outliers for RplV substitutions

Supplementary Table 3: List of RplV and RplD substitutions in NCBI AMR database

Supplementary Table 4: List of outliers for RplD substitutions

**Supplementary Table 3:**
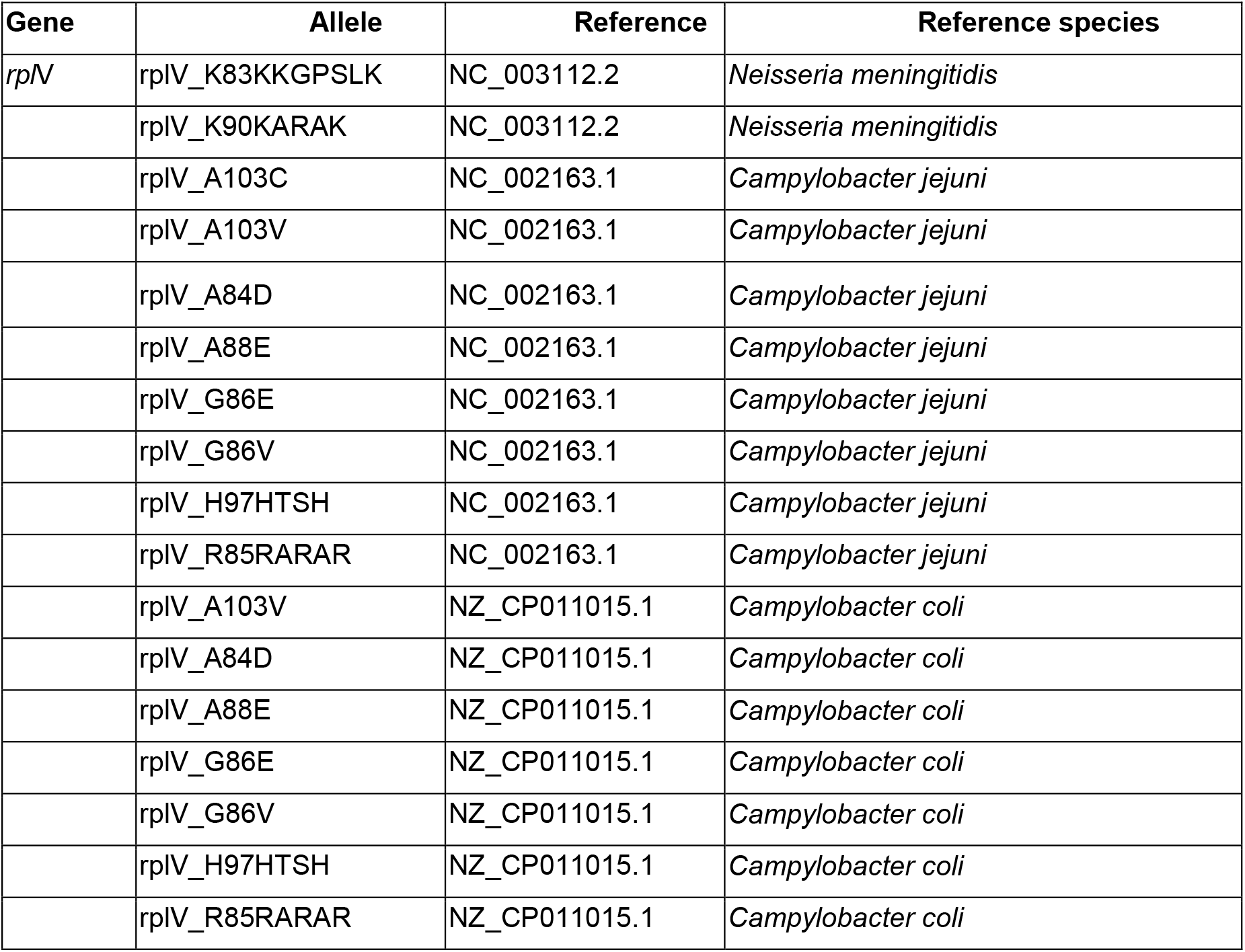
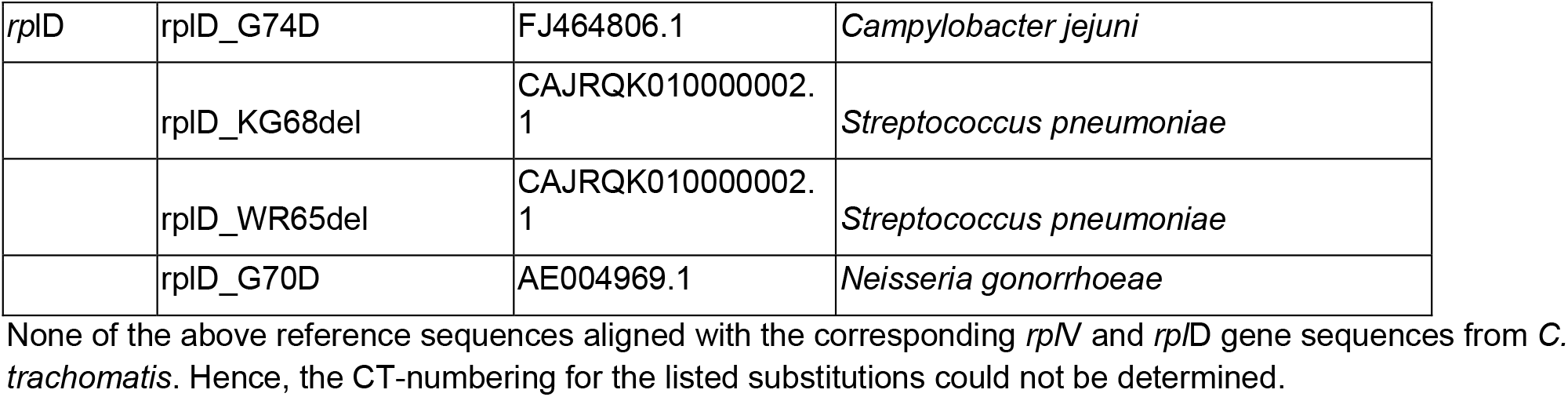
List of L22 (encoded by *rpl*V) and L4 (encoded by *rpl*D) substitutions in NCBI AMR database.

